# Acute inflammation sensitizes knee-innervating sensory neurons and decreases mouse digging behavior in a TRPV1-dependent manner

**DOI:** 10.1101/350637

**Authors:** Sampurna Chakrabarti, Luke A. Pattison, Kaajal Singhal, James R.F. Hockley, Gerard Callejo, Ewan St. John Smith

## Abstract

Ongoing, spontaneous pain is characteristic of inflammatory joint pain and reduces an individual’s quality of life. To understand the neural basis of inflammatory joint pain, we made a unilateral knee injection of complete Freund’s adjuvant (CFA) in mice, which reduced their natural digging behavior. We hypothesized that sensitization of knee-innervating dorsal root ganglion (DRG) neurons underlies this altered behavior. To test this hypothesis, we performed electrophysiological recordings on retrograde labelled knee-innervating primary DRG neuron cultures and measured their responses to a number of electrical and chemical stimuli. We found that 24-hours after CFA-induced knee inflammation, knee neurons show a decreased action potential generation threshold, as well as increased GABA and capsaicin sensitivity, but have unaltered acid sensitivity. The inflammation-induced sensitization of knee neurons persisted for 24-hours in culture, but was not observed after 48-hours in culture. Through immunohistochemistry, we showed that the increased knee neuron capsaicin sensitivity correlated with enhanced expression of the capsaicin receptor, transient receptor potential vanilloid 1 (TRPV1) in knee-innervating neurons of the CFA-injected side. We also observed an increase in the co-expression of TRPV1 with tropomyosin receptor kinase A (TrkA), which is the receptor for nerve growth factor (NGF), suggesting that NGF partially induces the increased TRPV1 expression. Lastly, we found that systemic administration of the TRPV1 antagonist A-425619 reversed the decrease in digging behavior induced by CFA injection, further confirming the role of TRPV1, expressed by knee neurons, in acute inflammatory joint pain.

## 1. Introduction

Pain is protective, but its dysregulation has a negative impact on an individual’s life, as well as a wider socioeconomic impact (St. John Smith, 2018; Vilen et al., 2017). Inflammatory joint pain is characteristic of many musculoskeletal disorders, often being more diffuse and longer lasting than cutaneous pain (Lewis, 1938). Numerous animal models of inflammatory joint pain are used experimentally (Gregory et al., 2013) and intra-articular complete Freund’s adjuvant (CFA) was chosen for use in this study because it produces robust, unilateral inflammation, alongside behavioral hyperalgesia (Keeble et al., 2005). Although spontaneous pain is a hallmark of joint pain, most rodent studies focus on evoked pain behaviors (Arendt-Nielsen, 2017; Schaible et al., 2002). Therefore, shifting the emphasis to study natural behaviors during painful conditions and their underlying molecular mechanisms might facilitate the translation of novel analgesics into clinics (Arendt-Nielsen, 2017).

A variety of techniques have been employed to better understand the molecular drivers of pain. For example, single cell RNA-sequencing studies have characterized dorsal root ganglia (DRG) neurons into distinct subpopulations (Hockley et al., 2018; Li et al., 2015; Usoskin et al., 2015), and electrophysiology and immunohistochemistry have shown that joint innervating DRG neurons possess distinct neurochemical and electrophysiological properties (da Silva Serra et al., 2016). These reports highlight the importance of specifically studying knee neurons, to fully understand the neural basis of joint pain.

Multiple ion channels are involved in transducing noxious stimuli; of these, the Transient Receptor Potential (TRP) channel family are critical to nociceptor chemosensitivity (Moore et al., 2018). Alongside endogenous mediators released during inflammation, e.g. protons, we have shown that articular neurons can be activated by the TRP channel agonists capsaicin (TRPV1), cinnamaldehyde (TRPA1) and menthol (TRPM8) (da Silva Serra et al., 2016). In relation to this, TRPV1^-/-^ and TRPA1^-/-^ mice show reduced pain behavior in CFA models of articular nociception (Chen et al., 2009; Fernandes et al., 2016). Although not tested after CFA-induced joint inflammation, TRPM8^-/-^ mice develop similar mechanical hyperalgesia to wildtype mice following CFA injection into the footpad (Caceres et al., 2017). Joint acidosis correlates with increased severity of inflammatory arthritis in humans, and acidic solutions activate and sensitize rodent nociceptors (Farr et al., 1985; Smith et al., 2011). Mice lacking the proton-gated ion channels acid-sensing ion channel 3 (ASIC3) and TRPV1 have reduced mechanical hyperalgesia following intra-articular CFA injection (Hsieh et al., 2017), but not following subcutaneous hind paw CFA injection, suggesting regional differences in pain processing (Staniland and McMahon, 2009). Subsequently, using magnetic resonance spectroscopic imaging, we found no evidence of tissue acidosis following subcutaneous CFA injection (Wright et al., 2018), suggesting absence of a key stimulus for ASIC3/TRPV1 activation, hence perhaps explaining the lack of phenotype observed in the hind paw CFA model. However, it remains unknown how proton-gated channels contribute to knee-innervating neuron activation during acute inflammation.

Regardless of the stimulus, once activated, sensory neuron input to the spinal cord is pre-synaptically modulated by GABA_A_ receptor signaling, the source of GABA being spinal cord interneurons (Sutherland et al., 2002). In adults, due to the high intracellular [Cl^-^] in DRG neurons compared to central nervous system neurons, activation of GABA_A_ receptors produces depolarization through Cl^-^ efflux (Chen et al., 2014). Furthermore, after subcutaneous CFA-induced inflammation in rat paws, the magnitude of GABA-evoked currents is increased in cutaneous DRG neurons (Zhu et al., 2012), as well as in isolated human DRG neurons incubated in an inflammatory soup (Zhang et al., 2015). However, the characteristics of GABA-evoked currents in knee-innervating neurons, and their modulation during inflammation are unknown.

Here, we hypothesized that sensitization of knee-innervating neurons following CFA-induced knee inflammation underlies changes in natural behavior.

## 2. Methods

### 2.1. Animals

All mice used in this study were 6-15 week old female C57BL/6J (Envigo); female mice were used because being female is a risk factor for inflammatory pain (Hannan, 1996). We did not control for estrus cycle because C57BL/6 mice show relatively stable behavior across the four stages of estrus, including pain behaviors (Meziane et al., 2007). Mice were conventionally housed in groups of 4-5 with nesting material and a red plastic shelter; the holding room was temperature controlled (21 °C) and mice were on a normal 12 hour/light dark cycle with food and water available *ad libitum.* This research was regulated under the Animals (Scientific Procedures) Act 1986 Amendment Regulations 2012 following ethical review by the University of Cambridge Animal Welfare and Ethical Review Body.

### 2.2. Knee joint intra-articular injections

Under anesthesia (ketamine, 100 mg/kg and xylazine, 10 mg/kg, i.p.) a single injection of the retrograde tracer Fast Blue (FB; 1.5 μl 2% in 0.9 % saline; Polysciences) was made intra-articularly through the patellar tendon into each knee to label knee-innervating neurons. For all experiments, anaesthetized mice were injected with 7.5 μl Complete Freund’s adjuvant (CFA; 10 mg/ml; Chondrex) intra-articularly through the patellar tendon in the left knee seven days after administration of FB. Knee width was measured with Vernier’s calipers before and 24-hours after CFA injection.

### 2.3. Digging behavior paradigm

Digging behavior testing was carried out in a subset of mice used for electrophysiological and immunohistochemistry studies. A standard 49 x 10 x 12 cm cage with a wire lid, filled with Aspen midi 8/20 wood chip bedding (LBS Biotechnology) tamped down to a depth of ~ 4 cm, was used as the test environment. Each mouse was tested individually using fresh bedding. Testing lasted three minutes and to avoid distractions food and water were not available. All digging experiments were carried out between 12:00 and 14:00 on weekdays in the presence of one male and one female experimenter.

#### 2.3.1. Testing protocol for assessing the impact of CFA-induced inflammation

Each day before assessment of digging, mice were habituated in the procedure room, in their home cage, for 30 min. On the training days (2 days before FB or CFA injection in the knee, Figure 1A) mice were allowed to dig twice, with a 30-minute break between sessions. All subsequent days were test days and, after habituation, mice were allowed to dig once. Test digs were recorded and the digging duration (time mice spent actively displacing bedding material with their front and hind limbs), as well as the number of visible burrows (crater-like sites in the cage with displaced bedding material) in test cages were counted immediately after the 3-minute test session. Digging duration was scored independently by the two experimenters by watching video recordings. Since the scores were well correlated (Pearson correlation R^2^ = 0.93), an average is reported.

**Figure 1:**
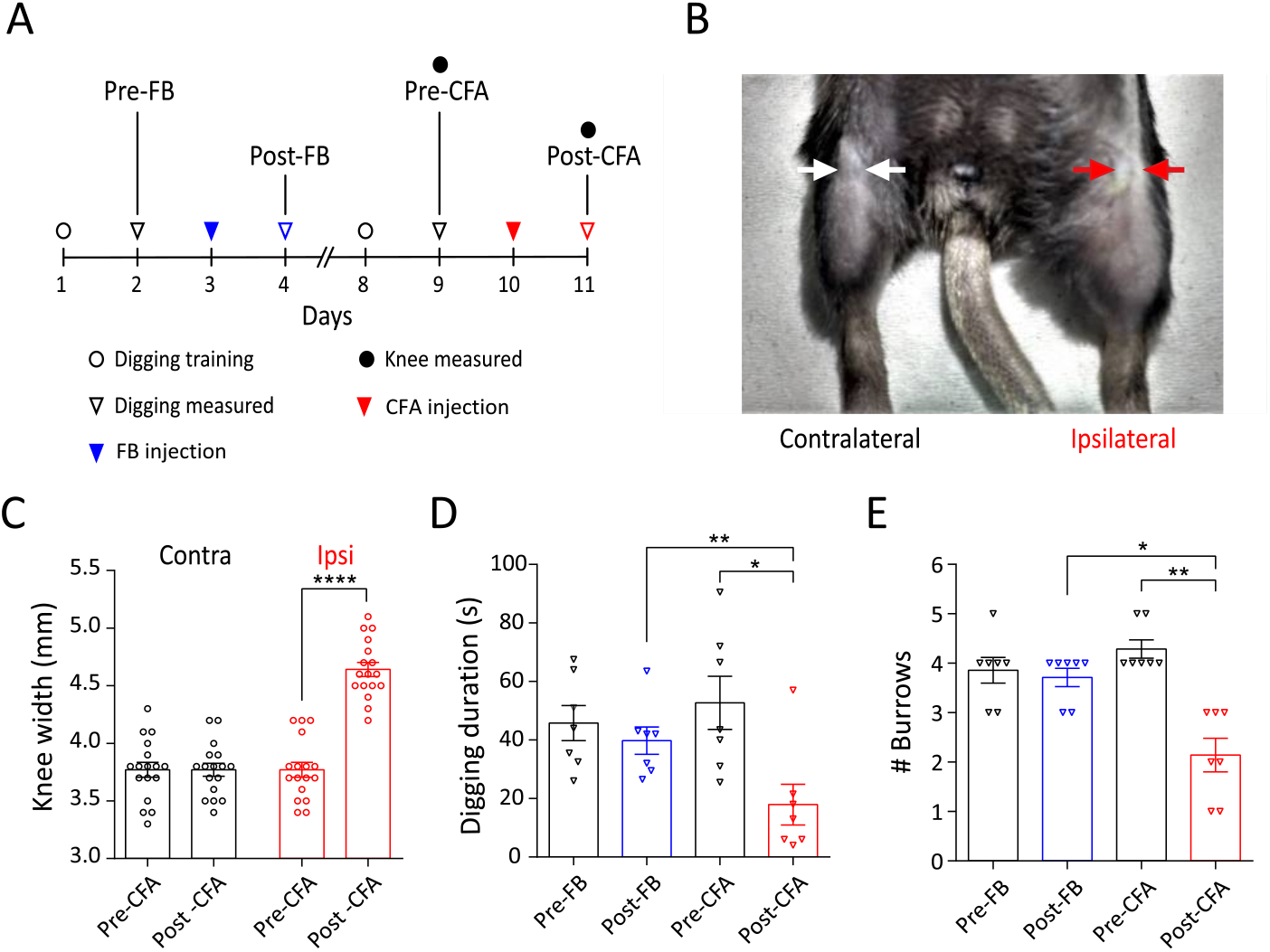
CFA model of acute knee inflammation in mice. A) Experimental timeline indicating when intra-articular injections, behavioral training and measurements were conducted. B) Representative picture of CFA injected (ipsilateral) and non-injected (contralateral) knee (shaved before injections). Arrows indicate where knee width was measured. C) Knee width on the day of CFA injection (pre) and 24-hours after CFA injection (post) in the ipsilateral (red dots) and contralateral (black dots) knee (n=17). **** indicates p < 0.0001, paired t-test. Plots of (D) time spent digging and (E) the number of visible burrows after a 3-minute digging test (n = 7). * indicates p < 0.05, ** indicates p < 0.01, repeated measures ANOVA, Holm-Sidak multiple comparison test. Error bars indicate SEM.

#### 2.3.2. Testing protocol for assessing the effect of TRPV1-antagonist, A-425619

Seventeen mice were trained as above and the day before CFA knee injections their digging behavior was measured twice with a 30 min interval in between. Since the digging duration (saline group: Run 1, 30.3 ± 7.1 s vs. Run 2, 33.8 ± 6.4 s, t(6) = 0.39, p = 0.7, n = 7; CFA group: Run 1, 32.2 ± 5.1 s vs. Run 2, 31.1 ± 6.1 s, t(9) = 0.21, p = 0.8, n = 10, paired t-test) and number of burrows (saline group: Run 1, 3.4 ± 0.4 vs. Run 2, 3.4 ± 0.4, t(6) = 0.0, p > 0.9, n = 7; CFA group, Run 1, 4.0 ± 0.3 vs. Run 2, 4.3 ± 0.4, t(9) = 1.1, p = 0.3, n = 10, paired t-test) did not change between these two test digs for either group, the measurement from the second run is reported as pre-CFA/saline. On the day of the injection, the mice were randomly split into two groups: the CFA group received CFA in one knee (as described above) and the control group received saline in one knee. 24-hours after the injections, digging activity was re-assessed, after which all mice received an intra-peritoneal dose of the TRPV1 antagonist, A-425619 (100 μmol/kg, Tocris, made up in 10% DMSO and 34% 2-hydroxylpropyl β-cyclodextrin, Sigma-Aldrich, in dH_2_O). Digging activity was measured again after 30 min as previous reports show maximal anti-nociceptive effect of A-425619 at this time point (Hillery et al., 2011). At the end of the study all the videos of test digs were independently assessed by the experimenters, who were blinded to the treatment conditions and stage of the experiment by an assistant, and an average of their scores is reported.

### 2.4. DRG neuron culture

Following cervical dislocation and decapitation, the spinal column was dissected and the lumbar (L2 to L5) DRG (that primarily innervate the knee joint) were collected (da Silva Serra et al., 2016; Rigaud et al., 2008). DRG from the CFA-injected and non-injected sides were separately collected in ice cold dissociation media containing L-15 Medium (1X) + GlutaMAX-l (Life Technologies), supplemented with 24 mM NaHCO_3_. DRG were then incubated in 3 ml type 1A collagenase (1 mg/ml with 6 mg/ml bovine serum albumin (BSA) in dissociation media; Sigma-Aldrich) for 15 min at 37 °C, followed by 30 min incubation in 3 ml trypsin solution (1 mg/ml with 6 mg/ml BSA in dissociation media; Sigma-Aldrich) at 37 °C. After removing the enzymes, the DRG were suspended in culture media containing L-15 Medium (1X) + GlutaMAX-l, 10 % (v/v) fetal bovine serum, 24 mM NaHCO_3_, 38 mM glucose, 2 % penicillin/streptomycin, dissociated by mechanical trituration with a 1 ml Gilson pipette and briefly centrifuged (160 g, 30 s; Biofuge primo, Heraeus Instruments; Hanau, Germany). Supernatants containing dissociated DRG neurons were then collected in a fresh tube. This was repeated five times. Finally, the cells were plated onto poly-D-lysine and laminin coated glass bottomed dishes (MatTek, P35GC-1.5-14-C) and incubated (37 °C, 5 % CO_2_) for 4-, 24- or 48-hours depending upon the experiment.

### 2.5. Immunohistochemistry

FB labelled and CFA-injected mice were transcardially perfused, firstly with PBS, followed by 4 % (w/v) paraformaldehyde (PFA; in PBS, pH 7.4) under terminal anesthesia (sodium pentobarbital; 200 mg/kg, i.p.). L2 – L5 DRG were collected from the CFA-injected and non-injected sides and post-fixed for 1-hour (4 % PFA), followed by an overnight incubation in 30 % (w/v) sucrose (in PBS) at 4 °C for cryoprotection. DRG were next embedded in Shandon M-1 Embedding Matrix (Thermo Fisher Scientific), snap frozen in 2-methylbutane (Honeywell International) on dry ice and stored at −80 °C. Embedded DRG were sectioned (12 μm) using a Leica Cryostat (CM3000; Nussloch, Germany), mounted on Superfrost Plus microscope slides (Thermo Fisher Scientific) and stored at −20 °C until staining. One to three sections were chosen at random for analysis from each CFA-injected and contralateral side of four mice.

Slides were defrosted, washed with PBS-tween and blocked in antibody diluent solution: 0.2 % (v/v) Triton X-100, 5 % (v/v) donkey serum and 1 % (v/v) bovine serum albumin in PBS for 1-hour at room temperature before overnight incubation at 4 °C with primary antibodies anti-TRPV1 (1:1000, rabbit polyclonal, Abcam ab31895) and anti-TrkA (1:1000, goat polyclonal, R&D systems AF1056) in antibody diluent. Slides were washed three times using PBS-tween and incubated with species-specific conjugated secondary antibodies (1:1000, anti-rabbit Alexa-488 (Invitrogen, A21206) and anti-goat Alexa-568 (Invitrogen, A11057)) for 2-hours at room temperature (20-22 °C). Slides were washed in PBS-tween, mounted and imaged with an Olympus BX51 microscope (Tokyo, Japan) and QImaging camera (Surrey, Canada). Exposure levels were kept constant for each slide and the same contrast enhancements were made to all slides. Negative controls without the primary antibody showed no staining with either secondary.

Using ImageJ, the mean gray value of each neuron in a DRG section was measured and normalized between the highest and lowest intensity neuron for that section. The threshold used for scoring a neuron as positive for a stain was set as the normalized minimum gray value across all sections + 2 times SD.

### 2.6. Whole cell patch-clamp electrophysiology

At least three DRG neurons from the CFA-injected side (CFA) and contralateral (Cntrl) side from each animal were recorded. To try and maintain an even distribution of animals across all time points at which recordings were made, the same culture was used for 4-hours, 24-hours and 48-hours wherever possible, however, due to the paucity of labelled neurons and variability of culture conditions, this was not possible in some cases. In total, each of the conditions contained DRG neurons from 6-8 mice. At each time point recordings were made for-4 hours. The extracellular solution contained (in mM): NaCl (140), KCl (4), MgCh_2_ (1), CaCh_2_ (2), glucose (4) and HEPES (10) adjusted to the required pH (> 6.0) with NaOH. For solutions with pH < 6.0, MES was used instead of HEPES. Patch pipettes of 4-9 MΩ were pulled with a P-97 Flaming/Brown puller (Sutter Instruments; Novato, CA, USA) from borosilicate glass capillaries and the intracellular solution used contained (in mM): KCl (110), NaCl (10), MgCh_2_ (1), EGTA (1), HEPES (10), Na_2_ATP (2), Na_2_GTP (0.5) adjusted to pH 7.3 with KOH. Recordings were made using a HEKA EPC-10 amplifier (Lambrecht, Germany) and the corresponding Patchmaster software. FB labelled neurons were identified by their fluorescence upon excitation with a 365nm LED (Cairn Research; Faversham, United Kingdom). For all experiments, DRG neurons were held at −60 mV and whole cell currents were acquired at 20 kHz. Only neurons where an action potential (AP) could be evoked in response to current injections and had a resting membrane potential more negative than −40 mV were analyzed. Images of neurons were captured using a 40x objective on a Nikon Eclipse Ti-S microscope and a Zyla 5.5 sCMOS camera (Andor; Belfast, United Kingdom), followed by pixel to μm conversion to calculate their diameter using ImageJ software.

### 2.6.1. Testing Protocols

#### Action potential generation

APs were generated by 80 ms current injections of 150-1050 pA in 50 pA steps (Figure 2A). If an AP was generated at 150 pA, the neuron was retested with current injections of 0–1000 pA, in 50 pA steps. Threshold, amplitude, half peak duration (HPD), afterhyperpolarization (AHP) duration and AHP amplitude (Figure 2A) were measured using Fitmaster software (HEKA) or IgorPro software (Wavemetrics) as described before (Bonin et al., 2013; Djouhri et al., 2001; Kim et al., 1998). AP amplitude is defined as the peak of the AP from the repolarized membrane potential; HPD is the AP width at half-amplitude, AHP was obtained by fitting the decay to a single exponential function and AHP amplitude was obtained by subtracting the peak voltage during AHP from the repolarized membrane potential (see Figure 2A for more detail).

**Figure 2:**
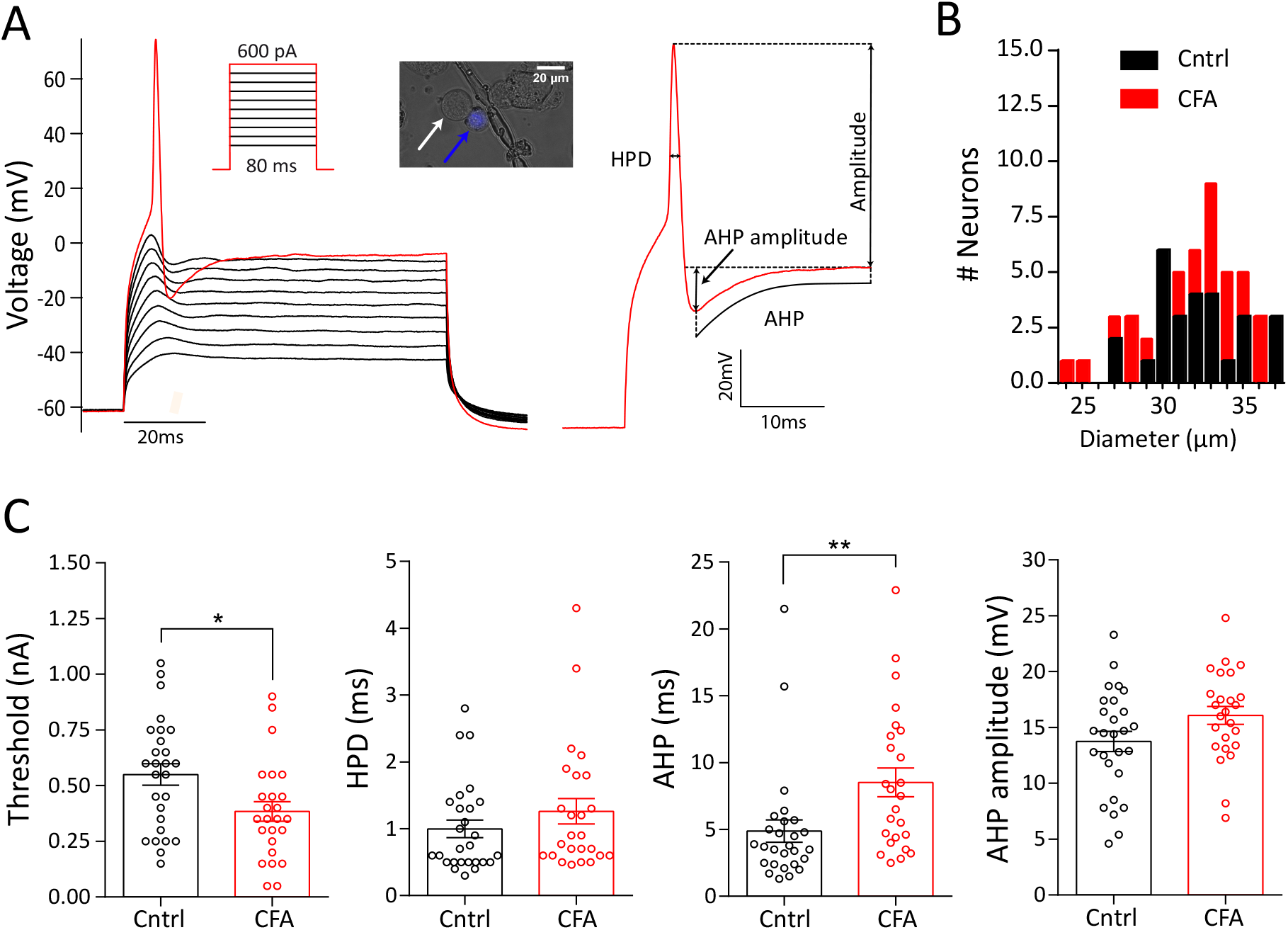
Action potential properties of knee neurons following acute knee inflammation. A) Action potential generation protocol (inset) and response along with a close-up of an evoked action potential from a knee neuron showing the different parameters measured. Example of a labeled knee neuron (blue arrow) and an unlabeled neuron (white arrow) shown in inset. Scale bar indicates 25 μm. B) Frequency distribution of neuronal diameter in μm of CFA (red, n = 25) and Cntrl (black, n = 27) neurons. C-F) Distribution of threshold (C), half peak duration (HPD) (D), afterhyperpolarization (AHP) (E) and AHP amplitude (F) after 4-hours in culture. Error bars indicate SEM, * indicates p < 0.05, ** indicates p < 0.01, unpaired t-test.

#### Acid sensitivity and TRP-agonist sensitivity

Solutions for determining acid sensitivity (pH 7, pH 6, pH 5) and TRP-agonist sensitivity (capsaicin, cinnamaldehyde and menthol) were applied in a random order through a gravity-driven 12 barrel perfusion system (Dittert et al., 2006) to DRG neurons in 5 s pulses with at least 30 s wash period (with pH 7.4) between stimuli. Solutions of 10 μM capsaicin (1 mM stock in 100% ethanol; Sigma-Aldrich), 100 μM cinnamaldehyde (10 mM stock in 100% ethanol; Merck) and 100 μM menthol (20 mM stock in 100% ethanol; Alfa Aesar) were made up in pH 7.4 extracellular solution from their respective stock solutions. Control experiments (n = 3) with ethanol did not evoke any currents from neurons. Current amplitude was measured in Fitmaster (HEKA) by subtracting the maximum peak response from the baseline (average of the first 3 s before stimulation), which was then normalized by dividing by neuron capacitance to give current density.

#### GABA sensitivity

100 μM GABA (100 mM stock in water; Sigma-Aldrich) was applied for 5 s followed by a 20 s wash with pH 7.4, then 100 μM GABA and 250 μM bicuculline (100 mM in DMSO; Sigma-Aldrich) were applied together for 5 s. Recovery from bicuculline block was assessed after 1 min by a subsequent 100 μM GABA application for 5 s. To test for the presence of the GABA_A_-δ subunit, 100 μM of the δ agonist 4,5,6,7-tetrahydroisoxazolo[5,4-c]pyridin-3-ol hydrochloride (THIP, 100 mM stock in distilled water; Sigma-Aldrich) was applied for 5 s.

### 2.7. Statistical Analysis

Comparison of CFA and Cntrl neuron electrophysiological properties was performed using a Student’s unpaired t-test (two-sided), while comparison across time (4-hours, 24-hours and 48-hours) was made using a one-way ANOVA followed by a Holm-Sidak multiple comparison test. To compare proportions in patch-clamp and immunostaining experiments, a chi-squared test was used. Knee width before and after inflammation with CFA was compared using a Student’s paired t-test (two-sided). Digging behavior of mice in different groups was compared using a repeated measures one-way ANOVA followed by a Holm-Sidak multiple comparison test. Data are presented as mean ± standard error of mean.

## 3. Results

### 3.1. Inflammation and reduced well-being in mice after unilateral knee joint CFA injection

Injection of CFA into the knee joint produces acute and chronic inflammatory pain in mice that models human arthritis (Gauldie et al., 2004). To produce acute inflammation, 7-days after labelling knee-innervating neurons by FB injection, CFA was injected into the left knee joint of mice (timeline in Figure 1A). After 24-hours, we observed robust swelling indicative of edema and inflammation of the ipsilateral knee (pre-CFA, 3.8 ± 0.07 mm vs. post-CFA, 4.6 ± 0.06 mm, t(16) = 22.2, p < 0.0001, n = 17, paired t-test), but not the contralateral knee (pre-CFA, 3.7 ± 0.07 mm vs. post-CFA, 3.7 ± 0.06 mm, n =17, t(16) = 0.0, p > 0.9, paired t-test, Figure 1B, 1C); a pilot study found that FB injection alone induced no change in knee width (n = 6, pre-FB, 3.9 ± 0.05 mm, post-FB, 4.0 ± 0.05 mm, t(5) = 0.89, p = 0.4, paired t-test)

Spontaneous pain is one of the most distressing features of arthritic conditions that reduces the overall feeling of well-being (Hawker et al., 2008; Treharne et al., 2005) and it has been suggested that burrowing and digging behaviors function as indicators of well-being in mice, such that a reduction in burrowing behavior during inflammatory pain indicates diminished well-being (Deacon, 2006; Jirkof, 2014). Based on this hypothesis, we measured digging behavior in a subset of mice (n = 7, timeline Figure 1A, see Video, Supplementary Digital Content 1, which demonstrates post-FB and post-CFA digging behavior of a mouse). During a 3-minute test period, neither the time spent digging (pre-FB, 45.8 ± 5.9 s vs. post-FB, 39.8 ± 4.7 s, F(1.8,11.3) = 9.8, t (6) = 1.1, ANOVA, Holm-Sidak’s multiple comparison test, Figure 1D), nor the number of burrows dug (pre-FB, 3.8 ± 0.3 vs. post FB, 3.7 ± 0.2, F(1.7,10.4) = 21.8, t (6) = 0.5, ANOVA, Holm-Sidak’s multiple comparison test, Figure 1E) was altered following FB injection. However, CFA injection resulted in decreased digging time (pre-CFA, 52.7 ± 9.1 s vs. post-CFA, 17.9 ± 6.9 s, F(1.8,11.3) = 9.8, t (6) = 3.9, p = 0.003, ANOVA, Holm-Sidak’s multiple comparison test, Figure 1D) and a decreased number of burrows dug (pre-CFA, 4.3 ± 0.2 vs. post-CFA, 2.1 ± 0.3, F(1.7,10.4) = 21.8, t (6)= 6.3, p = 0.0003, ANOVA, Holm-Sidak’s multiple comparison test, Figure 1E), suggesting a significant impact to well-being, elicited by joint inflammation. Comparison of post-FB and post-CFA time points also showed that CFA-injection produced a reduction in the time spent digging (F(1.8,11.3) = 9.8, t (6) = 4.8, p = 0.003 ANOVA, Holm-Sidak’s multiple comparison test) and in the number of burrows dug (F(1.7,10.4) = 21.8, t (6) = 4.3, p = 0.0003, ANOVA, Holm-Sidak’s multiple comparison test).

### 3.2. Action potential threshold of knee neurons decreases after intra-articular CFA injection

To determine if altered sensory input may underlie the inflammation-induced decrease in digging behavior observed, we characterized the electrical properties of knee-innervating DRG neurons isolated from mice undergoing unilateral CFA knee injection as above (CFA neurons); DRG neurons innervating the contralateral knee were used as control (Cntrl neurons). *In vivo* recordings from rat joint afferents have shown increased excitability during inflammation (Grubb et al., 1991). Therefore, we hypothesized that AP threshold would decrease in mouse knee neurons following CFA-injection.

We recorded electrically-evoked APs from CFA neurons (Figure 2A) (CFA, n = 25) and Cntrl neurons (n = 27) with similar diameters (t(50) = 0.47, p = 0.6, Figure 2B, Table 1) after 4-hours in culture. The AP threshold of CFA neurons was ~ 30 % lower compared to Cntrl neurons (t(50) = 2.52, p = 0.01, unpaired t-test) (Figure 2C, Table 1), suggesting hyperexcitability and sensitization of these neurons. By contrast, there was no change in the HPD (t(50) = 1.2, p = 0.3, unpaired t-test, Figure 2D), AHP amplitude (t(50) = 1.9, p = 0.06, Figure 2F) or amplitude (t(50) = 0.3, p = 0.7, unpaired t-test, Table 1) of the electrically-evoked APs following inflammation, whilst the AHP increased (t(50) = 2.7, p = 0.009, unpaired t-test, Figure 2E).

### 3.3. Acid-sensitivity does not change in knee neurons following inflammation

Tissue acidosis occurs in some forms of inflammation in humans (Farr et al., 1985; Fujii et al., 2015), and in rodents there are also contrasting findings (Andersson et al., 1999; Wright et al., 2018). We used a range of pH solutions to determine if knee neuron acid-sensitivity changes following knee CFA injection. Depending upon the pH stimulus used, virtually all neurons responded with an inward current (pH 7 = 80.6 ± 4.6 %, pH 6 = 98.0 ± 2.0 % and pH 5 = 100 %, combined percentage from CFA and Cntrl groups). A small proportion of neurons showed a rapidly activating transient inward current followed by a sustained phase (transient + sustained) at pH 6 and pH 5. These currents were likely mediated by acid-sensing ion channels (ASICs) and their frequency was unaltered in CFA neurons (CFA, 16.3 ± 3.7 %; Cntrl, 11.1 ± 3.7 %, combined mean of pH 6 and pH 5) (Figure 3A, 3B). Sustained currents were the dominant response at all the pH tested (Figure 3A, C) and no difference was observed in the amplitude of response to any pH stimulus between CFA and Cntrl neurons (Figure 3D-F, Table 2). Overall, this suggests that there was no significant change in acid sensitivity in articular neurons during acute inflammation.

**Figure 3:**
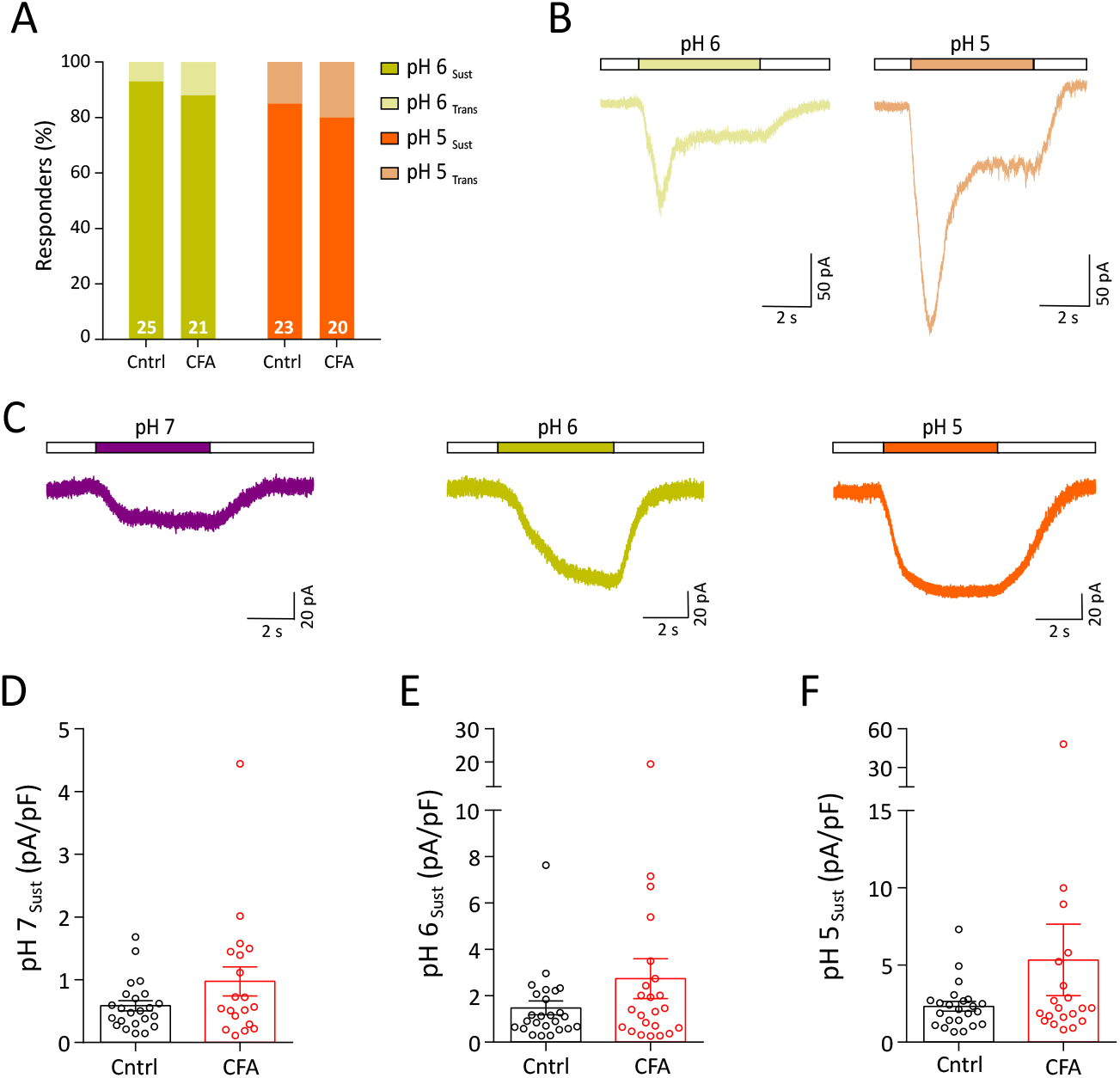
Acid sensitivity of knee neurons after acute inflammation. A) Percentage frequency of transient + sustained vs. sustained only response to pH 6 (transient + sustained, light yellow; sustained, dark yellow) and pH 5 (transient + sustained, light orange; sustained, orange) B) Example traces of transient + sustained acid currents in response to pH 6 (light yellow) and pH 5 (light orange). C) Example traces of sustained response to pH 7 (purple), pH 6 (dark yellow) and pH 5 (orange). D-F) Peak current density of sustained pH 7, pH 6 and pH 5 responses (CFA: red dots, Cntrl: black dots).

### 3.4. Inflammation increases the proportion of TRPV1 expressing knee neurons

Sensory neurons express a variety of TRP channels involved in thermo- and chemosensing and these are both activated and sensitized by a range of inflammatory mediators (Moore et al., 2018). To determine if any change in TRP-mediated chemosensitivity occurs in knee neurons following acute inflammation, we tested 10 μM capsaicin, 100 μM cinnamaldehyde and 100 μM menthol, agonists of TRPV1, TRPA1 and TRPM8 respectively.

The percentage of capsaicin-sensitive CFA neurons increased significantly compared to Cntrl neurons (CFA, 72 % vs. Cntrl, 44 %, X^2^(1) = 4.0, p = 0.04, chi-sq test, Figure 4A). However, there was no change in the peak current density of capsaicin-evoked currents (t(28) = 1.0, p = 0.3, unpaired t-test, Table 2). By contrast, no change in the percentage of cinnamaldehyde-sensitive (CFA, 40 %, Cntrl, 67 %, X^2^(1) = 3.7, p = 0.06; chi-sq test, Figure 4B) or menthol-sensitive neurons (CFA, 35 %, Cntrl, 51 %, X^2^(1) = 1.3, p = 0.2, chi-sq test, Figure 4C) was observed, nor was there any difference in the peak current density of either cinnamaldehyde-or menthol-evoked currents following inflammation (Table 2, cinnamaldehyde, t(26) = 0.6, p = 0.5; menthol, t(21) = 0.2, p = 0.9, unpaired t-test). An increased proportion of knee DRG neurons responding to capsaicin in inflammation suggests recruitment of a previously capsaicin “silent” population (Figure 4D).

**Figure 4:**
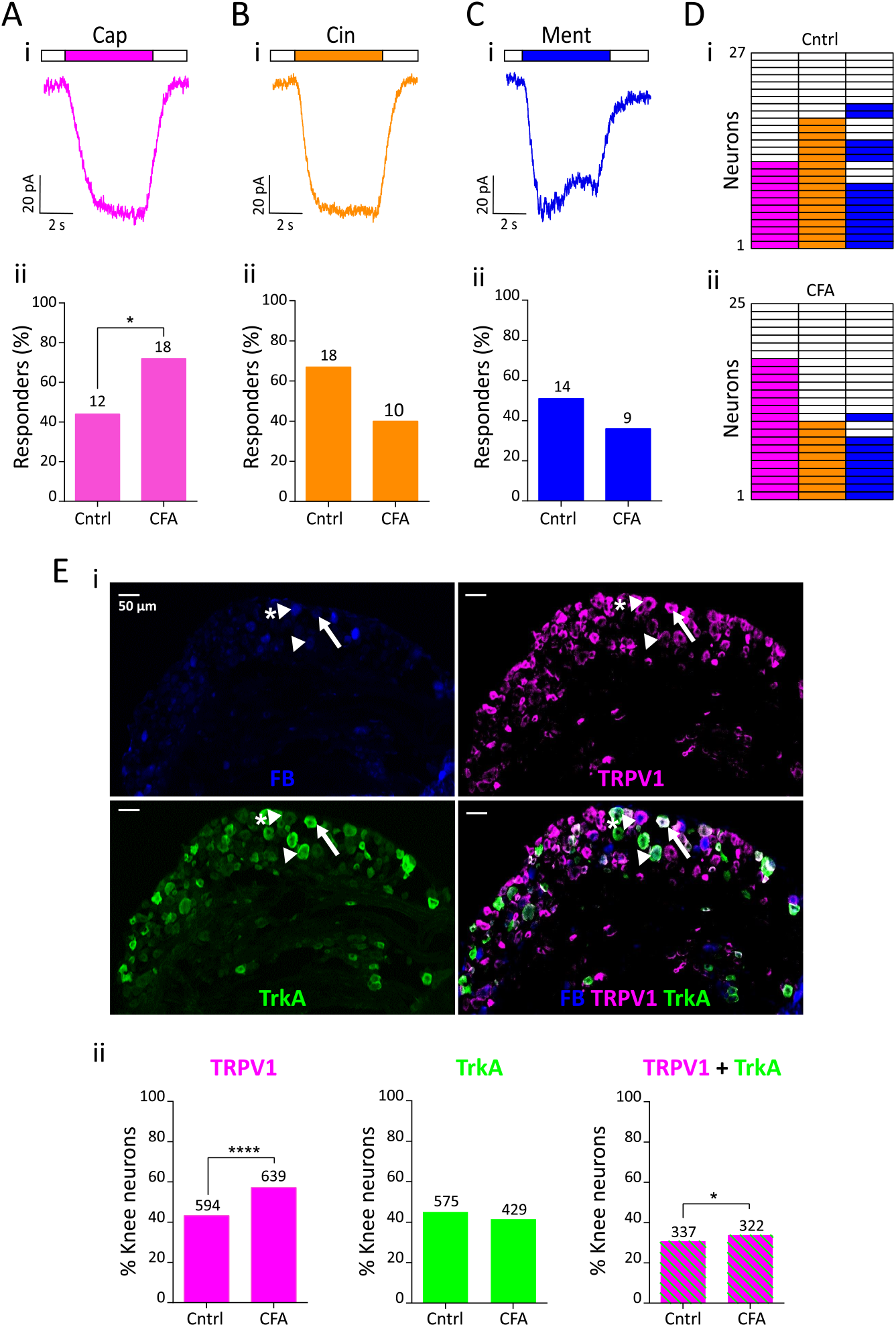
TRP agonist response profile of knee neurons following acute knee inflammation. Representative traces of 10 μM capsaicin (pink, Ai), 100 μM cinnamaldehyde (orange, Bi) and 100 μM menthol (blue, Ci) response from a CFA neuron and their respective percentage frequency (Aii, Bii, Cii). The numbers above the bars indicate the number of responsive neurons. D) Heat map of Cntrl (i, n = 27) and CFA (ii, n = 25) neurons responding to capsaicin, cinnamaldehyde and menthol. E (i) Representative images of a whole DRG section from L4 showing fast blue (FB) labeling from the knee (blue), TRPV1 expression (pink), TrkA expression (green) and a merged image; sections from an animal injected with CFA. White arrowhead shows a knee neuron that only expresses TrkA, white arrowhead with asterisk shows a knee neuron that expresses only TRPV1 and white arrow shows a knee neuron that co-expresses TRPV1 and TrkA. (ii) Proportion of knee neurons (L2-L5) that express TRPV 1 (pink), TrkA (green) and both TRPV 1 and TrkA (green and pink stripes) from Cntrl (n = 1334) and CFA (n = 1089) injected side. Numbers above the bars represent neurons stained positive with respective antibodies. * indicates p < 0.05, **** indicates p < 0.0001, chi-sq test.

To confirm that an increase in the proportion of capsaicin sensitive knee neurons is due to increased TRPV1 expression in knee neurons, we conducted immunohistochemistry on whole DRG sections (L2-L5) from ipsilateral (CFA, n = 1089 neurons) and contralateral (Cntrl, n = 1334 neurons) sides of four mice (Figure 4Ei). A significantly higher proportion of CFA neurons displayed TRPV1 expression (59.7%) compared to Cntrl neurons (44.6 %, X^2^(1) = 48.03, p < 0.0001, chi-sq test, Figure 4Eii). Nerve growth factor (NGF) is an important inflammatory mediator implicated in arthritis that binds to its high affinity receptor tropomyosin receptor kinase A (TrkA) and can increase TRPV1 expression (Ji et al., 2002). Hence, we hypothesized that the increased TRPV1 expression observed will occur in TrkA positive neurons, leading to increased TRPV1 and TrkA coexpression. Although, the proportion of knee neurons expressing TrkA (CFA, 39.3 %, Cntrl, 42.6 %) (Figure 4Eii) was not greater in CFA neurons than Cntrl neurons, there was a small, yet significant, increase in the proportion of CFA neurons co-expressing TRPV1 and TrkA compared to Cntrl neurons (CFA, 24.8 %; Cntrl, 29.8 %, X^2^(1) = 5.6, p = 0.02, chi-sq test, Figure 4Eii). Taken together, our data suggests that NGF-TrkA signaling contributes, alongside as yet unknown pathways, to driving increased TRPV1 expression following CFA.

### 3.5. Sustained GABAA-evoked current magnitude increases in knee neurons following inflammation

An inflammation-induced increase in amplitude of GABA-evoked currents has been reported in skin-labeled rat DRG neurons, suggesting that altered GABAergic signaling may be important in inflammatory pain (Zhu et al., 2012). Two types of GABA response were observed. In the first, the current reached a peak within 1.5 s of GABA application and was classified as transient + sustained (TS). Two subtypes of TS responses were observed: one showing relatively monophasic desensitization throughout GABA application (Figure 5Ai) and a second showing a far slower desensitization that produced a more stable sustained phase (Figure 5Aii); due to the paucity of neurons in the first subgroup and since no response returned to baseline within 5 s, both types were included in the TS group. The second type of GABA-evoked response did not reach its peak within the first 1.5 s, showed no desensitization and was classified as sustained (S, Figure 5Aiii). The frequency of TS and S responses in CFA and Cntrl neurons was similar (CFA: TS, 48 %, S, 28 % vs. Cntrl: TS, 52 %, S, 44 %, Figure 5 Bi). We also observed that 100 μM GABA evoked a response in 76 % of CFA and 96 % of Cntrl neurons (X^2^(1) = 4.6, p = 0.03, chi-sq test, Figure 5 Bi). Although no difference in GABA-evoked peak current density was observed overall between CFA and Cntrl neurons (Figure 5Bii, Table 2), GABA-evoked S type currents were of larger magnitude in CFA than Cntrl neurons (CFA, 8.0 ± 4.1 pA/pF, n =7 vs. Cntrl, 1.7 ± 0.6 pA/pF, n = 14, t(19) = 2.1, p = 0.04, unpaired t-test), whereas no such difference was observed for TS type currents (CFA, 6.7 ± 2.4 pA/pF, n =12 vs. Cntrl, 6.5 ± 1.9, n = 12, t(22) = 0.05, p = 0.9, unpaired t-test) (Figure 5Biii).

**Figure 5:**
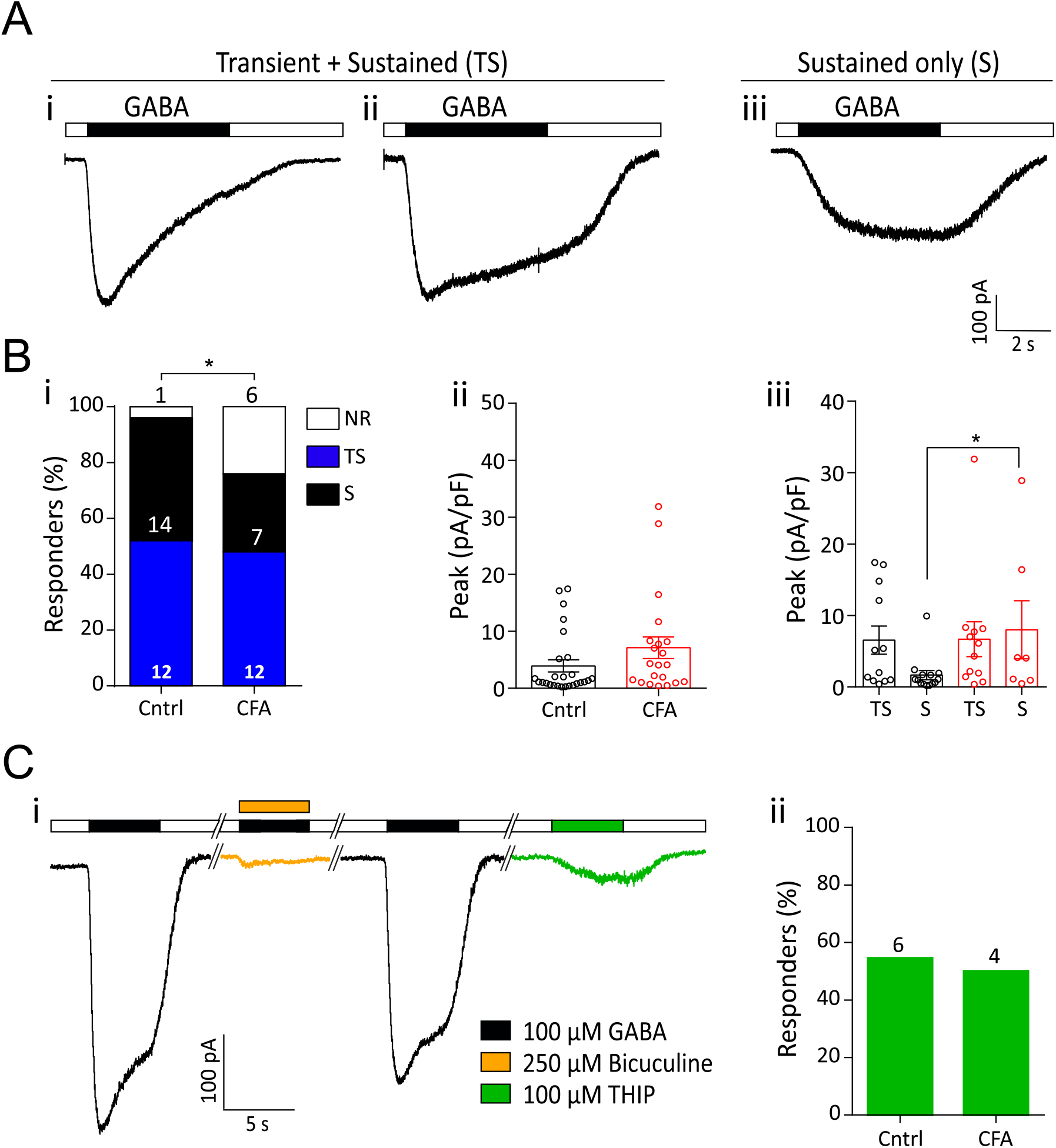
GABA-evoked currents from knee neurons after acute inflammation. A) Representative traces of 100 μM GABA-evoked currents showing both transient and sustained phases (i, ii) and currents with only a sustained phase (iii). B (i) Percentage frequency of GABA-evoked transient + sustained currents (blue bars), sustained currents (black bars) and non-responders (white bars) in Cntrl and CFA neurons. (ii) Peak current density of all GABA-evoked currents (Cntrl: n = 26, CFA: 19) and (iii) GABA-evoked currents split into TS and S type (Cntrl: black dots; CFA: red dots). TS = transient + sustained currents, S = sustained only currents, error bars indicate SEM, * indicates p < 0.05, unpaired t-test. C) i - Example trace of bicuculline block (yellow) of GABA-evoked currents and THIP-evoked current (green) in a single neuron. ii – percentage frequency of Cntrl (n = 11) and CFA (n = 8) neurons with a GABA response that also responded to THIP. The numbers above the bars indicate the number of responsive neurons.

Due to inflammation producing an increase in the magnitude of S type GABA-evoked currents, we investigated the subunit composition of these GABA receptors. The GABA_A_ antagonist, bicuculline, inhibited GABA responses by more than 50 % in 6/7 CFA DRG and 8/12 Cntrl DRG (Figure 5Ci), with all GABA-evoked currents being sensitive to bicuculline to some extent. These results concur with the previous observation that a fraction of GABA-evoked currents in rat DRG neurons is bicuculline insensitive (Lee et al., 2012). S type GABA_A_ currents can be mediated by δ subunit containing GABA_A_ receptors (Lee et al., 2012) and thus we tested the ability of the δ subunit containing GABA_A_ receptor agonist THIP to evoke currents in knee neurons. A sustained current in response to THIP was observed in 4/8 CFA and 6/11 Cntrl neurons, indicating expression of δ-subunit containing GABA_A_ receptors in knee neurons (Figure 5Ci, Cii). The peak current density of THIP-evoked currents was similar (t(8) = 0.18, p = 0.9, unpaired t-test) in CFA (0.6 ± 0.2 pA/pF) and Cntrl (0.5 ± 0.1 pA/pF) neurons. However, not all the neurons with S type GABA response showed THIP sensitivity. Combined with the presence of bicuculline insensitive currents, the lack of response to THIP in all neurons with S currents suggests expression of either *ρ* subunit containing GABA_A_ receptors, or perhaps GABA_B_ receptors, both of which produce sustained currents in response to GABA (Chebib and Johnston, 2001). Taken together, our data indicate the presence of diverse GABA_A_ receptor subunits in CFA and Cntrl neurons and demonstrate an increase in GABA-mediated neuronal excitation following knee inflammation.

### 3.6. Time-in-culture leads to loss of inflammation-induced sensitization of knee neurons

The period of time that neurons are left in culture for (time-in-culture) and constituents of the culture medium can greatly alter neuronal excitability (Scott and Edwards, 1980; Winter et al., 1988). To systematically characterize the effect of time-in-culture in our system, we repeated the experiments described above on the CFA and Cntrl knee neurons after 24-hours (CFA, n = 25; Cntrl, n = 22) and 48-hours (CFA, n = 23; Cntrl, n = 22) in culture.

No change was observed in the resting membrane potential, capacitance or AP amplitude (Table 1) across the three time points. By contrast, whereas the AP threshold was similar across time points for CFA knee neurons (F(2,70) = 2.4, p = 0.09, ANOVA), it decreased for Cntrl neurons after 48-hours in culture compared to 24-hours (F(2,68) = 3.4, t(68) = 2.6, p = 0.03, ANOVA, Holm-Sidak multiple comparison test, Table 1). Interestingly, the inflammation-induced decrease in AP threshold of CFA neurons compared to Cntrl neurons that was observed at 4-hours (Figure 2C) and 24-hours (t(45) = 3.3, p = 0.002, unpaired t-test, Table 1) had disappeared after 48-hours in culture (Table 1, t(43) = 0.8, p = 0.44, unpaired t-test), which suggests that time-in-culture leads to a loss of inflammation-induced hyperexcitability. There was also a distinct AP widening characterized by an increase in both HPD and AHP in the 48-hours group for Cntrl neurons, but just the HPD in CFA neurons. HPD increased by ~ 60 % after 48-hours in both CFA (F(2,70) = 2.4, t(70) = 2.3, p = 0.01, ANOVA, Holm-Sidak multiple comparison test) and Cntrl (F(2,68) = 8.1, t(68) = 4.0, p = 0.007, ANOVA, Holm-Sidak multiple comparison test) neurons compared to the 4-hours group (Table 1). In Cntrl neurons, AHP increased by 191 % in the 48-hours group compared to 4-hours and 81 % compared to 24-hours group (F(2,68) = 10.0, p = 0.0002, ANOVA, Holm-Sidak multiple comparison test, Table 1). One CFA and one Cntrl neuron in the 48-hours culture group were excluded from this analysis because of a highly different AP waveform that could not be analyzed using the same measurements.

With regards to knee neuron acid sensitivity, the percentage of neurons with transient + sustained currents in response to pH 6 and pH 5 at 24-hours and 48-hours (see Figure, Supplementary Digital Content 2A showing time-in-culture effects on acid of knee neurons after inflammation) was similar to that at 4-hours (Figure 3A). Although a largely reversible increase in pH 7- (CFA and Cntrl) and pH 5-evoked (Cntrl) current amplitude was observed at 24-hours (pH 7: CFA, F(2,50) = 5.7, p = 0.005; Cntrl, F(2,52) = 9.3, p = 0.0003; pH 6: CFA, F(2,65) = 2.8, p = 0.06; Cntrl, F(2,60) = 2.6, p = 0.08; and pH 5: CFA, F(2,57) = 2.8, p = 0.06; Cntrl, F(2,54) = 5.2, p = 0.008, ANOVA) (Table 2), there was no clear trend across different pH stimuli for either CFA or Cntrl neurons.

For knee neuron TRP-mediated chemosensitivity, the CFA-induced increase in the proportion of capsaicin-sensitive neurons observed at 4-hours was maintained at 24-hours (CFA, 88 % vs. Cntrl, 50%, X^2^(1) = 8.1, p = 0.004, chi-sq test), but was absent in the 48-hours group (CFA, 54.2 % vs. Cntrl, 43.5 %, X^2^(1) = 0.5, p = 0.5, chi-sq test), a result that further confirms our observations that after 48-hours in culture the inflammation-induced effects are lost. In addition, the proportion of cinnamaldehyde (CFA, 33.3 %; Cntrl, 21.7 %) and menthol-sensitive (CFA, 33.3 %; Cntrl, 13.0 %) neurons after 48-hours in culture decreased compared to the other two time points, although this decrease was only significant in Cntrl neurons (cinnamaldehyde, X^2^(1) = 11. 9, p = 0.0006; menthol, X^2^(1) = 7.8, p = 0.005, chi-sq test) (see Figure, Supplementary Digital Content 2B, showing time-in-culture effects on TRP agonist of knee neurons after inflammation). No significant difference in the peak current density of TRP agonist-evoked currents was observed across time-in-culture (Table 2).

When GABA-evoked currents were split into TS and S types, no changes in the peak current density were observed between CFA and Cntrl after 24-hours or 48-hours in culture. However, combining both TS and S currents, CFA neurons had larger amplitude GABA-evoked currents than Cntrl neurons after 24-hours in culture (t(39) = 2.5, p = 0.01, unpaired t-test, Table 2); an effect that was not seen in the 48-hours group (t(32) = 1.2, p = 0.2, unpaired t-test, Table 2). Previously, it has been shown in cultured rat DRG neurons that the proportion of S type GABA-evoked currents increased with time-in-culture (Lee et al., 2012), a result replicated here in mouse knee neurons (CFA, X^2^(1) = 4.8, p = 0.03, Cntrl, X^2^(1) = 5.4, p = 0.02, chi-sq test, see Figure, Supplementary Digital Content 2C, showing time-in-culture effects on GABA sensitivity of knee neurons after inflammation). From this data, we would recommend not using DRG neurons for longer than 28-hours in culture to prevent loss of inflammation-induced sensitization events.

### 3.7. TRPV1 inhibition reverses inflammation induced decrease in digging behavior

From our characterization of CFA and Cntrl neurons, knee neuron TRPV1 expression emerged as an important target in CFA-induced knee inflammation. We hypothesized that TRPV1 underlies our observed knee inflammation-induced decrease in digging behavior observed in mice and that the TRPV1 antagonist A-425619 would thus reverse this decrease. On day 2 of our experimental timeline (Figure 6A), both the CFA group (n = 10) and saline group (n = 7) spent a similar amount of time digging (CFA, 31.1 ± 6.1 s; Saline, 33.8 ± 6.4 s) and produced a similar number of burrows (CFA, 4.3 ± 0.4; Saline, 3.4 ± 0.4) during a 3 min test in the digging cage. Following CFA/saline injection, the CFA group showed an expected decrease in digging duration (17.4 ± 5.2 s, F(1.8,16.6) = 10.1, t(9) = 3.6, p = 0.001, repeated measures ANOVA, Holm-Sidak multiple comparison test, Figure 6B), which was reversed 30 min after the injection of A-425619 (36.4 ± 7.0 s, F(1.8,16.6) = 10.1, t(9) = 3.9, p = 0.001, ANOVA, Holm-Sidak multiple comparison test, Figure 6B). A similar result was also obtained with the number of burrows (post-CFA, 2.1 ± 0.3, F(1.9,17.9) = 18.9, t(9) = 5.0; post-Ant, 4.0 ± 0.2, F(1.9,17.9) = 18.9, t(9) = 5.0, p < 0.0001, repeated measures ANOVA, Holm-Sidak multiple comparison test, Figure 6C). In contrast, the saline group did not show a decrease in digging behavior 24-hours after saline injection into the knee (Digging duration, 33.1 ± 5.3 s, F(1.5,9.0) = 0.6, t(6) = 0.1, p = 0.5; # Burrows, 3.7 ± 0.5, F(1.7,10.0) = 0.3, t(6) = 0.6, p = 0.7, repeated measures ANOVA, Holm-Sidak multiple comparison test, Figure 6D, 6E), further confirming that intra-articular injections do not affect digging behavior. The TRPV1 antagonist A-425619 also produced no effect on digging behavior in the saline group (Digging duration, 25.9 ± 6.3 s, F(1.5,9.0) = 0.6, t(6) = 0.8, p = 0.5; # Burrows, 3.3 ± 0.4, F(1.7,10.0) = 0.3, t(6) = 0.6, p = 0.7, ANOVA, Figure 6D, 6E), i.e. inhibiting TRPV1 does not interfere with spontaneous digging activity of mice. These results confirm that TRPV1 is an important mediator of inflammatory knee pain and contributes to the inflammation-induced decrease in digging behavior in mice.

**Figure 6:**
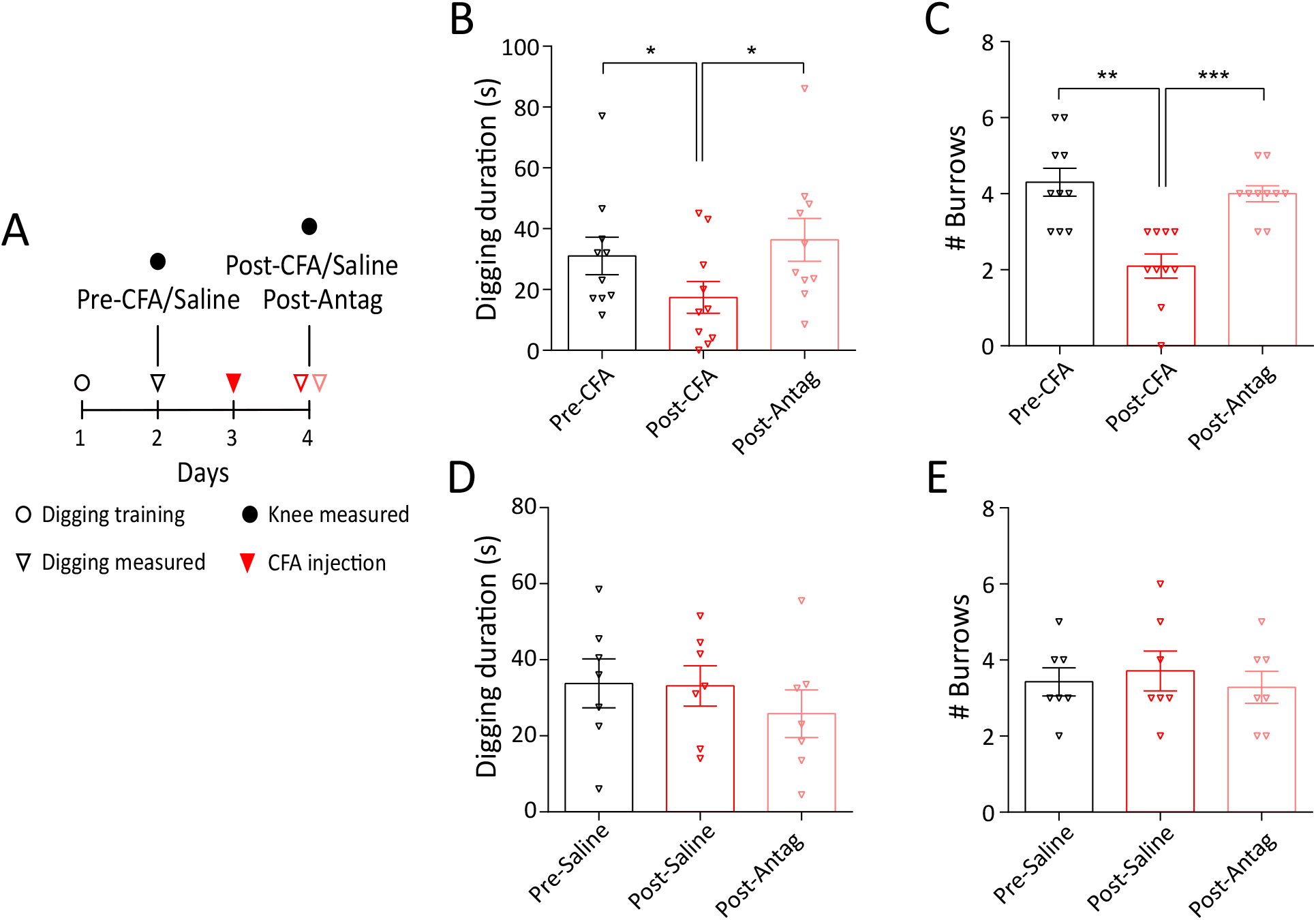
Digging behavior after injection of the TRPV1 antagonist A-425619. A) Experimental timeline highlighting the days where behavioral testing and injection was performed. Distribution of time spent digging and the number of burrows in the CFA (B, C, n = 10) and saline group (D, E, n = 7). * indicates p < 0.05, ** indicates p < 0.01, *** indicates p < 0.0001. rmANOVA, Holm-Sidak multiple comparison test. Error bars indicate SEM.

## 4. Discussion

CFA evokes both acute (within 18 hours) and chronic (> 6 weeks) inflammatory pain (Hsieh et al., 2017; Keeble et al., 2005), as demonstrated by altered evoked-pain behaviors, such as hot/cold plate tests. However, such evoked-pain tests are neither ethologically relevant to rodents, nor model experience of human chronic pain (Rice et al., 2008). Consequently, innate rodent behaviors, such as burrowing, digging and nest-building that seek to mirror human activities of daily living have been explored as pain assays (Deacon, 2006; Jirkof, 2014; Shepherd et al., 2018). A decrease in burrowing behavior has been used to assess pain-like behavior after CFA-induced knee inflammation in rats (Rutten et al., 2014) and digging is a similar innate behavior that is quick and easy to test in a laboratory (Deacon, 2006). Here we show that 24-hours after CFA-induced knee inflammation, mice spent significantly decreased time digging, suggesting that digging can be used as an ethologically relevant pain assay in mice; importantly, intra-articular injection of the retrograde tracer FB did not alter digging behavior.

To study the neural basis for the inflammation-induced change in behavior observed, we combined electrophysiology with retrograde labelling of the knee joints, i.e. those neurons exposed to the inflammatory insult. We found that knee neurons have a decreased AP threshold and increased AHP (although no change in AHP amplitude) following CFA-induced knee inflammation. The recorded increased AHP in CFA neurons is perhaps contradictory because an increase in current mediated by hyperpolarization-activated cyclic nucleotide-gated ion channels (HCN), and thus a decrease in AHP, is associated with repetitive firing, as shown by one study in trigeminal ganglia neurons after CFA-induced inflammation (Flake and Gold, 2005). However, other studies using CFA-induced hind limb inflammation either observed no change in AHP (like our data at 24-hours in culture) (Djouhri and Lawson, 1999) or reported a decrease in AHP only in C-fibers (Weng et al., 2012). Therefore, our data possibly demonstrates the multifaceted nature of the molecular mechanisms regulating AHP, a suggestion supported by the wide range of AHP values reported (Djouhri et al., 1998; Lawson et al., 1996).

TRP channels are cellular sensors of thermal, physical and chemical stimuli (Moore et al., 2018) and inflammatory mediators released during joint pain can sensitize TRP channels (Zhang et al., 2013). TRPV1 (Zhang et al., 2005), TRPA1 (Story et al., 2003) and TRPM8 (Behrendt et al., 2004) are expressed in nociceptors and are involved in hyperalgesia. However, the role of TRPV1 and TRPA1 in acute inflammatory joint pain is unclear. Some studies report that TRPV1^-/-^ mice do not develop CFA-(Keeble et al., 2005) or carrageenan- (Davis et al., 2000) induced hyperalgesia and others have shown that in DRG neurons TRPV1 protein expression increases in CFA-induced inflammation (Amaya et al., 2003; Ji et al., 2002; Yu et al., 2008). In direct contrast, other studies failed to show increased neuronal TRPV1 protein (Bär et al., 2004; Zhou et al., 2003) or mRNA expression (Ji et al., 2002) following CFA-induced inflammation. Similarly, TRPA1^-/-^ mice show normal development of CFA-induced hyperalgesia in some studies (Fernandes et al., 2011; Petrus et al., 2007), whereas others report attenuation of CFA- and carrageenan-induced hyperalgesia in TRPA1^-/-^ mice (Garrison and Stucky, 2014; Moilanen et al., 2012) or following administration of a TRPA1 antagonist (Bonet et al., 2013; Samer R Eid et al., 2008). We provide electrophysiological evidence that the proportion of capsaicin-sensitive knee neurons increases during acute inflammation from ~ 50% to ~80%, suggesting recruitment of a previously silent neuronal population. We validate this electrophysiology result through immunohistochemistry, demonstrating a similar magnitude increase in the proportion of TRPV1 expressing knee neurons; the observed TRPV1 expression in contralateral knee-innervating neurons reported here agrees with previous findings in rodents (Cho and Valtschanoff, 2008). Following that, we were able to reverse the inflammation-induced reduction in digging behavior in mice by systemic administration of the TRPV1 antagonist, A-425619, which suggests that the increased TRPV1 expression observed at the cellular level drives the pain behaviors. Multiple mechanisms may contribute to the increase in TRPV1 expression, for example, NGF released from a variety of cell types in arthritic joints may bind to TrkA and get retrogradely transported to DRG neurons to drive TRPV1 expression. Indeed, increased levels of NGF are found in rheumatic diseases and anti-NGF/TrkA therapy reduces arthritic pain in human and animal models (reviewed in (Seidel et al., 2013)). With regard to CFA-induced pain, it has been previously shown that the proportion of TrkA-expressing neurons innervating bone does not increase following CFA-induced bone pain (Sara Nencini et al., 2017). We also observed no change in TrkA expression, but did observe a slight, yet significant, increase in TrkA-TRPV1 co-expression following knee inflammation which suggests that NGF-TrkA signaling plays a role in the increased TRPV1 expression observed here.

Protons are algogens detected by sensory neurons through a variety of means, including ASICs, TRPV1, background K+ channels, chloride channels and proton-sensing G-protein coupled receptors (Holzer, 2009). There is conflicting evidence surrounding the role of ASICs in inflammatory pain, some studies showing no change, others decreased pain and still others increased pain (Price et al., 2001; Sluka et al., 2003). In terms of joint inflammation, ASIC3^-/-^, TDAG8^-/-^ (a proton-sensing GPCR) and TRPV1^-/-^ mice showed decreased hyperalgesia in an ankle inflammation model (Hsieh et al., 2017). In addition, Ikeuchi *et al* (Ikeuchi et al., 2009) observed upregulation of ASIC3 immunoreactivity in knee-innervating DRG neurons after carrageenan-induced acute knee inflammation in mice. By contrast, ASIC1, ASIC2 and ASIC3 knockout mice did not display attenuated thermal or mechanical hyperalgesia after intraplantar CFA injection, suggesting that these ASIC subunits play no role in cutaneous CFA-induced hyperalgesia (Staniland and McMahon, 2009). We evaluated direct proton activation of knee neurons and found that acid-evoked currents are neither enhanced, nor is there any change in the proportion of ASIC-like currents, following CFA-induced knee inflammation, results that might explain the lack of phenotype observed in certain studies.

Information transfer from peripheral to central afferents is modulated by GABAergic signaling. Although an inhibitory neurotransmitter in the central nervous system, GABA produces membrane depolarization through Cl^-^ efflux in DRG neurons owing to the relatively high intracellular [Cl^-^] (Sung et al., 2000). GABA-evoked currents in sensory neurons are predominantly GABA_A_ receptor mediated, (identified by their sensitivity to GABA_A_ antagonist, bicuculline) with diverse subunit composition (Lee et al., 2012). As shown previously in rat DRG neurons (Lee et al., 2012), we recorded both transient and sustained GABA-evoked currents from mouse knee neurons that were largely inhibited by the GABA_A_ antagonist bicuculline. In addition, our finding that approximately half of all knee neurons were activated by the GABA_A_-δ subunit agonist, THIP, suggests common expression of the δ-subunit, although the proportion of THIP-activated neurons did not change after CFA injection. The fact that bicuculline did not completely abolish GABA-evoked currents indicates the likely presence of *ρ* subunits in GABA_A_ receptors, as has previously been reported in rat DRG neurons (Lee et al., 2012), or the presence of GABA_B_ receptor expression (both *ρ* containing GABA_A_ receptors and GABA_B_ receptors are insensitive to bicuculline block). Under normal conditions, GABA-mediated excitation of primary afferent neurons causes primary afferent depolarization, an inhibitory action due to inducing inactivation of voltage gated sodium channels (i.e. depolarizing block). However, following tissue injury, primary afferent depolarization is larger and can actually generate AP to produce pain by excitation of presynaptic GABA_A_ receptors (Cervero et al., 2003; Willis, 1999). In this study, we found a decrease in number of GABA responsive CFA neurons, but an increase in the magnitude of GABA-evoked currents compared to Cntrl neurons indicating a potential GABA_A_ mediated decrease in presynaptic inhibition and an increase in presynaptic excitation.

Furthermore, we addressed how time-in-culture affects the electrophysiological properties of neurons isolated from mice undergoing CFA-induced inflammation. Acutely dissociated DRG neurons have been extensively studied with patch clamp electrophysiology, however the definition of “acute” varies. While some studies have recorded within 8-hours of dissection, it is more common to record after overnight incubation (18-28-hours). This variability can potentially confound results on inflammation-induced sensitization, as AP properties change with time-in-culture. Here we show AP broadening, and a slight decrease in AP threshold in neurons from the contralateral side as reported before during a similar time course (Scott and Edwards, 1980). We also show that time-in-culture does not affect inflammation-induced lowering of AP threshold up to 28-hours in culture, but that the lowered threshold is lost at 48-hours in culture. Understanding how time-in-culture alters neuronal excitability is important for correctly designing studies. For example, we (present study) and others (Lee et al., 2012) have shown that S type GABA-evoked responses increase in prevalence with time-in-culture. The present study also shows a decline in the frequency of response to capsaicin, cinnamaldehyde and menthol and the inflammation-induced increase in frequency of capsaicin-sensitive neurons, although maintained for up to 28-hours in culture is lost after 48-hours. By contrast, acid-sensitivity remained relatively stable across all time points.

In summary, we show that following unilateral acute inflammation of the knee, knee neurons have 1) a lower AP threshold 2) increased sensitivity to capsaicin and expression of TRPV1 and 3) an increasing trend in the magnitude of GABA-evoked currents up to 28-hours in culture. This specific sensitization signature of knee neurons highlights the importance of studying DRG neurons as a spatially heterogeneous group. Among these different modes of sensitization that likely contribute to the decreased mouse digging behavior following knee inflammation, an increase in TRPV1 function is perhaps most clinically relevant to inflammatory arthritis (reviewed in (Fernandes et al., 2013). Indeed, it has recently been shown that intra-articular administration of a TRPV1 antagonist attenuated early osteoarthritic pain in a mono-sodium iodoacetate model of osteoarthritis (Haywood et al., 2018). Historically in pre-clinical models, TRPV1 antagonists attenuated thermal hyperalgesia, but their efficacy in mechanical hyperalgesia and ongoing pain is unclear (Brandt et al., 2012). The peripherally restricted TRPV1 antagonist, A-425619, has been previously shown to reverse thermal and mechanical hypersensitivity in rat CFA models and also restore weight bearing ability in rat osteoarthritis models (Honore et al., 2005). Here we report that systemic administration of A-425619 is also able to restore the reduction in spontaneous digging behavior in mice brought about by CFA-induced knee inflammation, which further establishes TRPV1 as an important drug target in inflammatory joint pain.

## Acknowledgments

The authors would like to thank Julie Gautrey and Chris Cardinal for technical assistance with animal work. This work was supported by an Arthritis Research UK Grant (G.C. and E.St.J.S., RG 20930), a Rosetrees Postdoctoral Grant (J.R.F.H. and E.St.J.S., A1296), S.C. was supported by a Gates Cambridge Trust scholarship and L.A.P. was supported by the University of Cambridge BBSRC Doctoral Training Programme (BB/M011194/1).

## Data Statement

The datasets presented in this study can be accessed from the University of Cambridge Apollo data repository (https://doi.org/10.17863/CAM.24492).

---

Supplementary Digital Content 1: Video that shows a mouse digging after FB injection in the knee (left) and after CFA injection in the knee (right).mp4

**Table 1:**
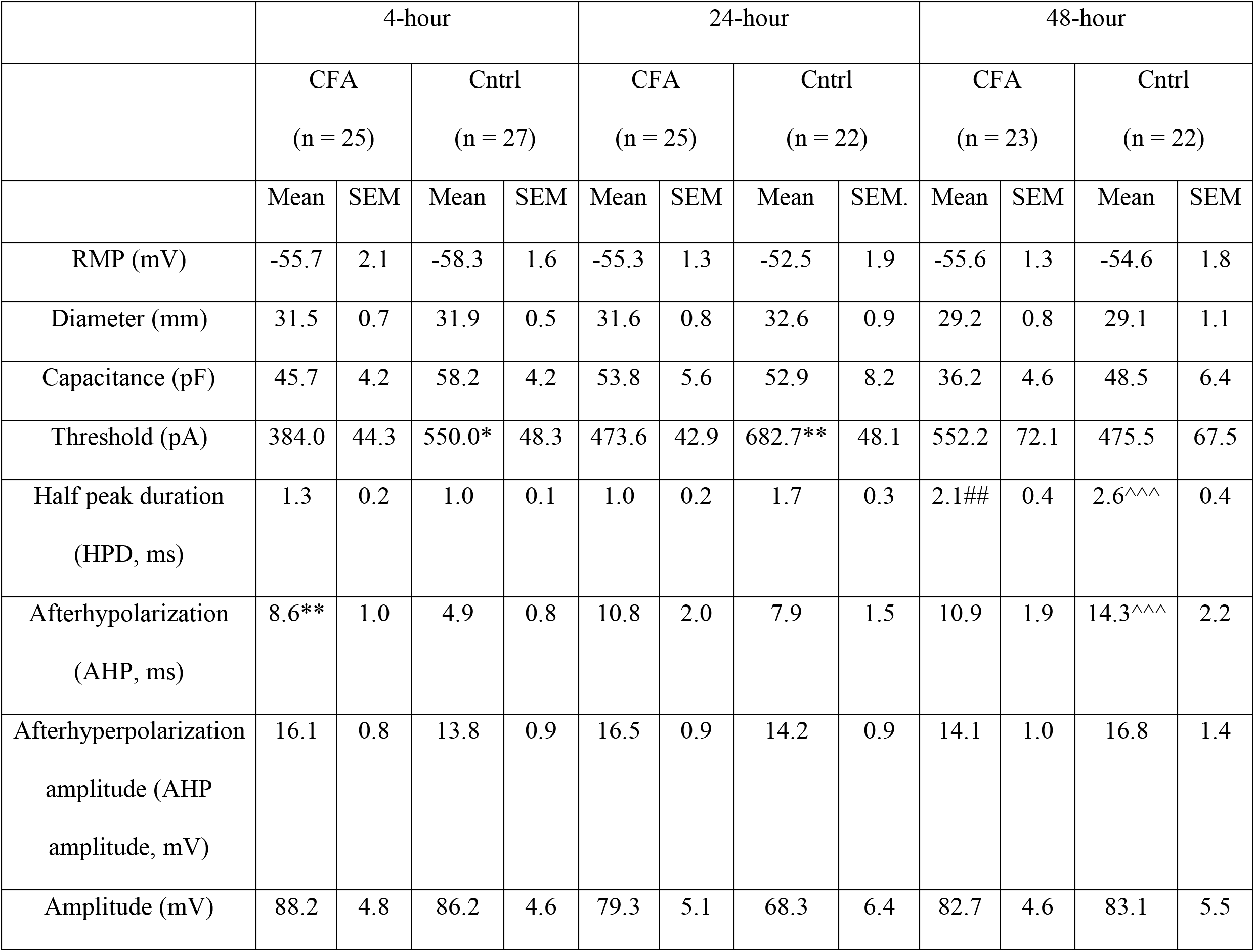
Properties of knee-innervating dorsal root ganglion neurons from the CFA injected side (CFA) and contralateral side (Cntrl) across time-in-culture. * = significance test of CFA vs Cntrl neurons in the same time group by unpaired t-test, # = significance test of CFA neurons amongst the three time points by ANOVA, ^ = significance test of Cntrl neurons amongst the three time points by ANOVA. */#/^ indicates p < 0.05, **/##/^^^ indicates p < 0.01, ***/###/^^^ indicates p < 0.0001.

**Table 2:** Sustained peak current densities of CFA and Cntrl DRG neurons in response to acid, TRP channel agonists (capsaisin, cinnamaldehyde and menthol) and GABA across time-in-culture. # = significance test of CFA neurons amongst the three time points by ANOVA, ^ = significance test of Cntrl neurons amongst the three time points by ANOVA. #/^ indicates p < 0.05, ##/^^^ indicates p < 0.01, ###/^^^ indicates p < 0.0001.

| Treatment | 4-hour |  |  |  |  |  | 24-hour |  |  |  |  |  | 48-hour |  |  |  |  |  |
| --- | --- | --- | --- | --- | --- | --- | --- | --- | --- | --- | --- | --- | --- | --- | --- | --- | --- | --- |
|  | CFA |  |  | Cntrl |  |  | CFA |  |  | Cntrl |  |  | CFA |  |  | Cntrl |  |  |
|  | Mean | SEM | n | Mean | SEM. | n | Mean | SEM | n | Mean | SEM | n | Mean | SEM | n | Mean | SEM | n |
| pH 7 | 1.0 | 0.2 | 19 | 0.6 | 0.08 | 23 | 4.0## | 1.0 | 19 | 5.7^^^ | 1.5 | 17 | 1.5 | 0.5 | 14 | 1.5 | 2.7 | 15 |
| pH 6 | 2.6 | 0.9 | 21 | 1.5 | 0.3 | 25 | 4.0 | 0.7 | 20 | 5.3 | 1.6 | 19 | 5.8 | 1.4 | 22 | 4.9 | 1.9 | 19 |
| pH 5 | 5.3 | 2.3 | 20 | 2.3 | 0.3 | 23 | 14.9 | 4.3 | 21 | 14.5^^ | 4.2 | 19 | 7.4 | 1.4 | 19 | 7.9 | 2.9 | 15 |
| Capsaicin | 4.4 | 2.8 | 18 | 0.8 | 0.1 | 12 | 1.8 | 0.5 | 22 | 4.6 | 2.1 | 11 | 10.3 | 5.1 | 13 | 6.4 | 4.0 | 10 |
| Cinnamaldehyde | 0.8 | 0.2 | 10 | 0.7 | 0.09 | 18 | 2.2 | 0.4 | 19 | 2.3 | 0.7 | 14 | 0.9 | 0.3 | 8 | 2.5 | 2.0 | 5 |
| Menthol | 0.8 | 0.2 | 9 | 0.9 | 0.2 | 14 | 1.0 | 0.2 | 12 | 1.6 | 0.4 | 9 | 1.2 | 0.7 | 8 | 2.8 | 2.2 | 3 |
| GABA | 7.2 | 2.0 | 19 | 3.9 | 1.1 | 26 | 11.8* | 2.6 | 22 | 4.3 | 0.9 | 19 | 8.3 | 2.7 | 18 | 4.3 | 1.7 | 16 |

**Supplementary Digital Content 2:**
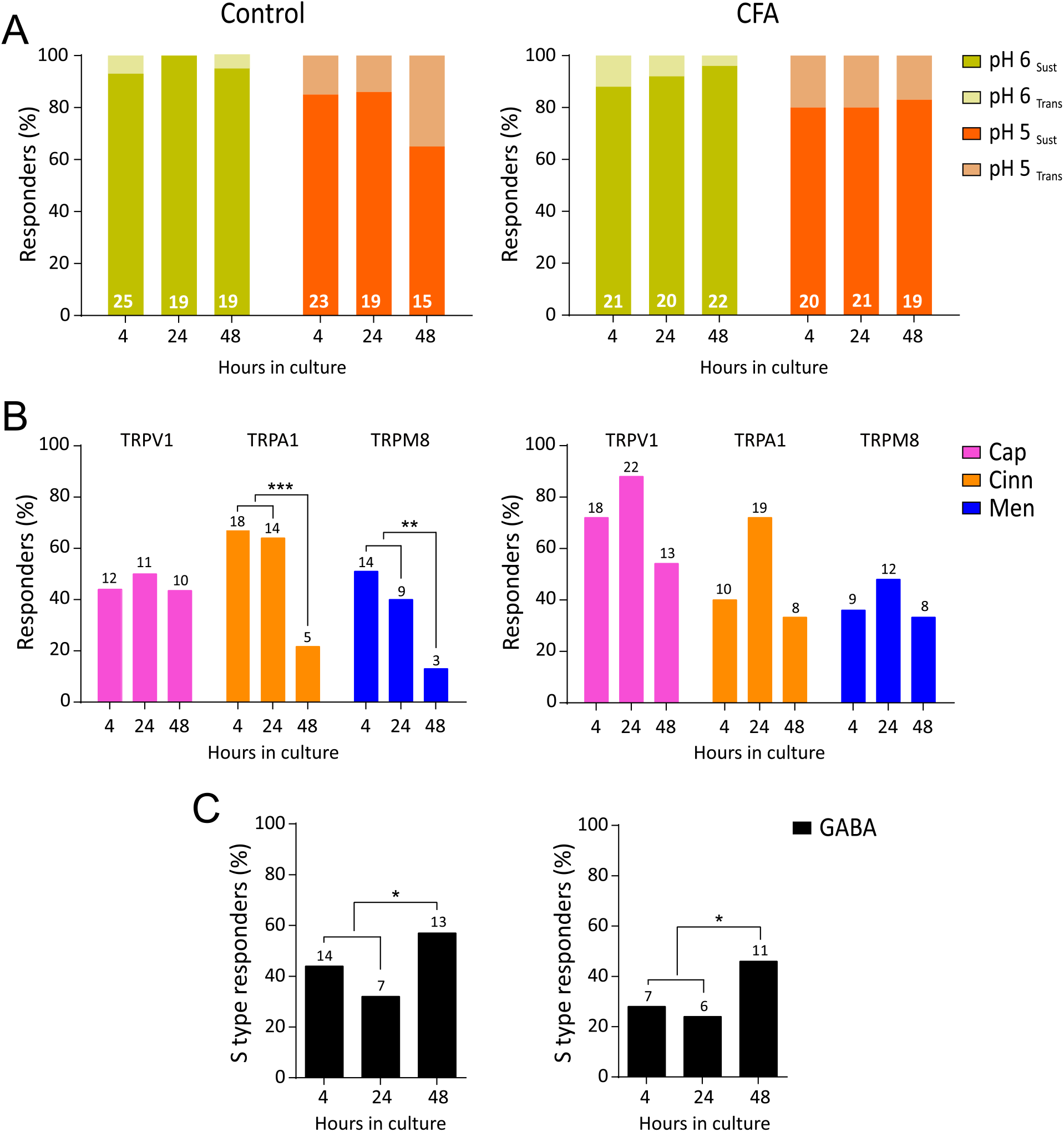
Figure showing time-in-culture effects on acid, TRP agonist and GABA sensitivity of knee neurons after inflammation.pdf. A) Proportion of Cntrl (left) and CFA (right) neurons responding to pH 6 (yellow) and pH 5 (orange) with a transient + sustained type current (light shade) and sustained only type current (dark shade) across 4, 24 and 48-hours. B) Percentage frequency of Cntrl (left) and CFA (right) neurons sensitive to capsaicin (pink), cinnamaldehyde (orange) and menthol (blue) across 4, 24 and 48-hours in culture. C) Percentage of Cntrl (left) and CFA (right) neurons that had a sustained GABA-evoked current across 4, 24 and 48-hours in culture. * indicates p < 0.05, chi-sq test. The numbers above the bars indicate the number of responsive neurons.

